# Theory of axo-axonic inhibition

**DOI:** 10.1101/2024.12.19.629419

**Authors:** Romain Brette

**Author notes:** Corresponding author: Romain Brette.

## Abstract

The axon initial segment of principal cells of the cortex and hippocampus is contacted by GABAergic interneurons called chandelier cells. The anatomy, as well as alterations in neurological diseases such as epilepsy, suggest that chandelier cells exert an important inhibitory control on action potential initiation. However, their functional role remains unclear, including whether their effect is indeed inhibitory or excitatory. One reason is that there is a relative gap in electrophysiological theory about the electrical effect of axo-axonic synapses. This contribution uses resistive coupling theory, a simplification of cable theory based on the observation that the small initial segment is resistively coupled to the large cell body acting as a current sink, to fill this gap. The main theoretical finding is that a synaptic input at the proximal axon shifts the action potential threshold by an amount equal to the product of synaptic conductance, driving force at threshold, and axial axonal resistance between the soma and either the synapse or of the middle of the initial segment, whichever is closer. The theory produces quantitative estimates useful to interpret experimental observations, and supports the idea that axo-axonic cells can potentially exert powerful inhibitory control on action potential initiation.

**Significance statement:** Chandelier cells form GABAergic synapses on the initial segment of pyramidal cells of the cortex and hippocampus. Despite their striking morphology and alterations in neurological diseases such as epilepsy, their functional role remains unclear. This study develops a quantitative theory to precisely assess the electrical impact of a synaptic input at the proximal axon. It shows that axo-axonic inhibition acts by shifting the action potential threshold proportionally to the synaptic conductance. This work underlines the role of chandelier cells in controlling action potential initiation and provides a quantitative tool to interpret experimental observations.

## Introduction

In most vertebrate neurons, action potentials (APs) initiate in a small axonal structure near the soma, the axon initial segment (AIS) (Bender and Trussell, 2012; Coombs et al., 1957). In principal cells of the cortex and hippocampus, the AIS harbors GABA_A_ receptors contacted by interneurons, most of which are axo-axonic cells (AACs), also called “chandelier” cells (about 60% of synapses on the proximal axon of visual cortical cells of mice (Schneider-Mizell et al., 2021)). Chandelier cells are electrically coupled fast-spiking inhibitory neurons (Howard et al., 2005; Woodruff et al., 2011) with a distinct morphology, forming cartridges along the AIS of principal cells (DeFelipe et al., 1985; Fairén and Valverde, 1980; Somogyi, 1977).

This striking anatomy suggests that AACs exert control on AP initiation, and may have an important role in normal function, or in the maintenance of the excitatory-inhibitory balance. *In vivo*, AACs targeting principal cells of the hippocampus fire at the peak of theta oscillations, when principal cells are least active, and they stop firing during sharp waves (Klausberger et al., 2003; Viney et al., 2013). AACs targeting principal cells of the primary visual cortex respond strongly to locomotion and visuomotor mismatch (Seignette et al., 2024). Alterations in axoaxonic inhibition have been reported in several neuropathologies, but their causal implication is unknown (DeFelipe, 1999; Gallo et al., 2020; Vivien et al., 2023; Wang et al., 2024).

Despite these suggestive clues, the precise effect of axoaxonic inhibition on neural function remains unclear. In fact, there has been some controversy over whether it is indeed inhibitory or excitatory, as a number of *in vitro* studies indicate that chandelier cells have a depolarizing effect (Szabadics et al., 2006; Woodruff et al., 2009, 2010), while *in vivo* they have an inhibitory action on hippocampal pyramidal cells of adult mice (Dudok et al., 2021). This discrepancy might be related to developmental changes in the reversal potential of chloride, the ionic carrier of GABA_A_ currents (Pan-Vazquez et al., 2020; Rinetti-Vargas et al., 2017). However, a recent *in vitro* study indicates that axoaxonic synapses still reduce excitability even when the reversal potential is substantially depolarized (Lipkin and Bender, 2023). *In vivo*, blocking AAC activity globally does not produce obvious effects (Jung et al., 2023). The interplay of axoaxonic synaptic currents with structural plasticity of the AIS is also unclear. In cultures, activity can induce a distal displacement of the AIS (Grubb et al., 2011; Jamann et al., 2018), but the axoaxonic synapses stay in place (Muir and Kittler, 2014; Pan-Vazquez et al., 2020; Wefelmeyer et al., 2015). The electrophysiological significance of this fact is not entirely obvious.

What makes interpretation difficult is the relative lack of theory about axonal inhibition. Indeed, classical excitability theory has focused on isopotential and spatially homogeneous situations (Hodgkin and Huxley, 1952; Izhikevich, 2006), while theoretical work on synaptic integration has addressed dendrites, notably by Rall (Rall, 2011). Here I provide a biophysical understanding of the effect of AIS synapses on excitability, based on resistive coupling theory, a simplification of cable theory that applies to the particular geometrical configuration where APs initiate in a thin axon close to a much larger cell body, a situation typical of many vertebrate neurons (Brette, 2013; Goethals and Brette, 2020; Kole and Brette, 2018). The theory results in a simple finding: axoaxonic inhibition raises the somatic threshold for AP initiation in proportion of synaptic conductance, axosomatic coupling resistance and driving force at threshold: Δ*V*^*^ = *g*_*s*_*R*_*a*_(*E*_*GABA*_ − *V*^*^) (where V* is threshold). Axoaxonic inhibition is most powerful on the second half on the AIS, and is more stable when it does not move along with the AIS.

## Results

### Resistive coupling between soma and proximal axon

In most vertebrate neurons, APs initiate on a thin axon, close to a much larger cell body. In this situation, the soma acts as a current sink for transmembrane currents in the proximal axon (Fig. 1A). Indeed, the conductance towards the soma is much larger than towards the distal axon or through the axonal membrane. Thus, as is shown in Figure 1B in a simple biophysical model (dendrite, soma and axon of diameter 6 µm, 30 µm and 1 µm, respectively), a current injected at the proximal axon produces essentially the same voltage response at the soma than a current injected directly at the soma (grey, axonal injection 40 µm from the soma; black: somatic injection).

**Figure 1.**
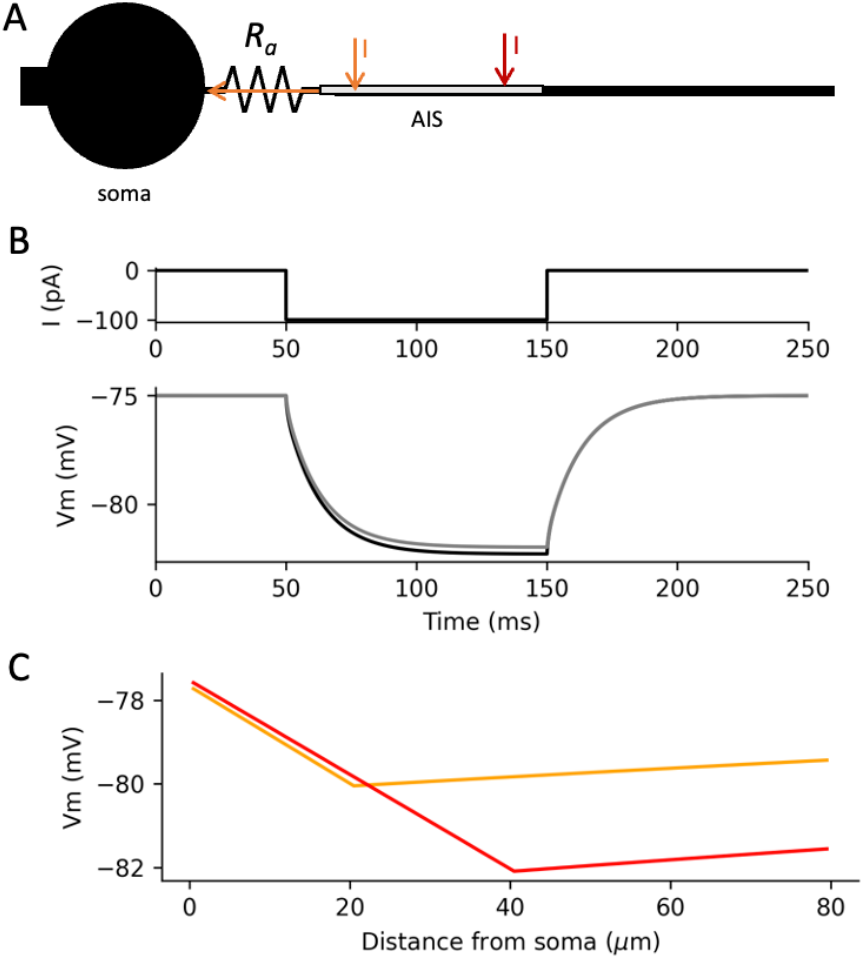
Resistive coupling between soma and proximal axon, in a simple biophysical model. A, A current I injected in the proximal soma (orange, 20 µm; red, 40 µm) flows mostly resistively towards the cell body. B, The somatic membrane potential in response to a current (top) injected in the soma (bottom, black) is essentially the same as when the current is injected in the proximal axon (grey, 40 µm). C, This results in a linear (ohmic) change in membrane potential between soma and injection site, with slope proportional to I (here shown at t = 5 ms).

It follows that a current injected in the proximal axon flows mostly longitudinally towards the soma, and therefore resistively, thereby producing a nearly instantaneous ohmic voltage gradient between the soma and the current injection site (Fig. 1C). This gradient follows Ohm’s law, and is therefore equal to *R*_*a*_.*I*, where *R*_*a*_ is the axial (longitudinal) resistance between the two sites, on the order of 1 MΩ/µm in layer 5 pyramidal cells of the cortex. Thus, with a cylindrical axon, the membrane potential varies linearly between soma and injection site, with a slope proportional to *I*. It follows that the same current injected 40 µm away from the soma (red) produces a local hyperpolarization that is twice larger than the same current injected 20 µm away (orange).

This phenomenon can be observed in dual soma-axon patch-clamp recordings of layer 5 cortical pyramidal cells, as shown in Figure 2 where the axonal electrode is placed 75 µm away from the soma (data reanalyzed from (Hu and Bean, 2018)). When a -50 pA current pulse is injected at the soma, it hyperpolarizes both the soma and the axon in the same way (left column), by about 5 mV. When the same current is injected in the axon, a ∼5 mV hyperpolarization also results in the soma, but an additional hyperpolarization is seen at the axonal site (∼4 mV), which closely tracks the step current (right column). A zoom on the response onset shows that the establishment of this gradient is very fast, as expected from a resistive phenomenon (inset). The amplitude of the soma-axon voltage gradient matches the theoretical expectation based on Ohm’s law (50 pA x 75 µm × 1 MΩ/µm = 3.75 mV).

**Figure 2.**
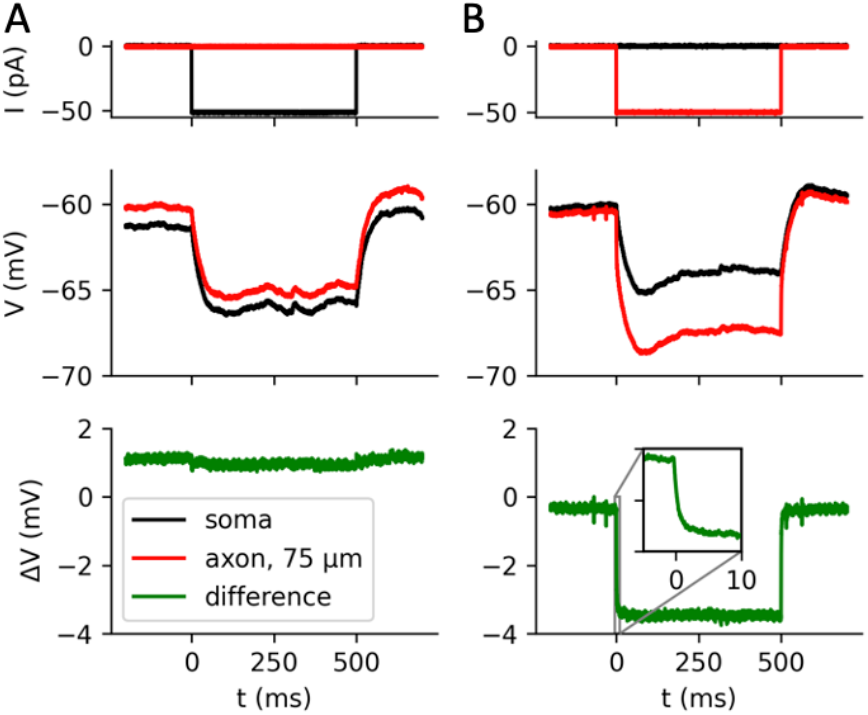
Resistive coupling in simultaneous patch-clamp recordings of soma and axonal bleb of pyramidal cortical neurons (data from (Hu and Bean, 2018)). A, A negative current injected at the soma (top) results in the same hyperpolarization at the soma (middle, black) and proximal axon (red, 75 µm). The bottom trace shows the difference (green). B, The same current injected at the proximal axon (top), hyperpolarizes the axon (middle, red) more than the soma (black), with an additional ohmic component (green) due to the resistance between the soma and axonal injection site.

Thus, the soma and proximal axon are resistively coupled. This simple resistive coupling greatly simplifies theoretical analysis. It allows quantitative understanding of the different factors influencing the initiation of APs (Brette, 2013; Goethals and Brette, 2020; Telenczuk et al., 2017), their backpropagation to the soma (Goethals et al., 2021; Hamada et al., 2016), and their extracellular signature (Teleńczuk et al., 2018). Although relatively simple, the theory can be somewhat counter-intuitive. For example, a prediction of the theory is that increasing the axial resistance between the soma and AIS, as when the AIS shifts distally, makes the cell more excitable (Brette, 2013), and this prediction has been experimentally confirmed by pinching the proximal axon of pyramidal cells with glass pipettes (Fékété et al., 2021).

We now examine the impact of synaptic currents at the proximal axon on AP initiation.

### Modulation of action potential initiation by axonal synaptic currents

A synaptic current on the proximal axon flows into the soma. Therefore, its effect on synaptic integration is essentially the same as a somatic current. However, there is an additional effect on AP initiation, as shown in Figure 3A on a simple biophysical model (see Methods, Numerical simulations). In this simulation, a hyperpolarizing current is injected in the middle of the AIS, extending from 5 to 35 µm from the soma, and an action potential is triggered by a depolarizing current pulse at the soma (1 nA, 5 ms). The somatic phase plot (dV/dt vs. V) shows the characteristic bimodal shape, where the first mode corresponds to the AP transmitted by the AIS and the second mode corresponds to the AP regenerated at the soma. The AP threshold (inflexion point of the first mode) increases with increasing synaptic current intensity.

**Figure 3.**
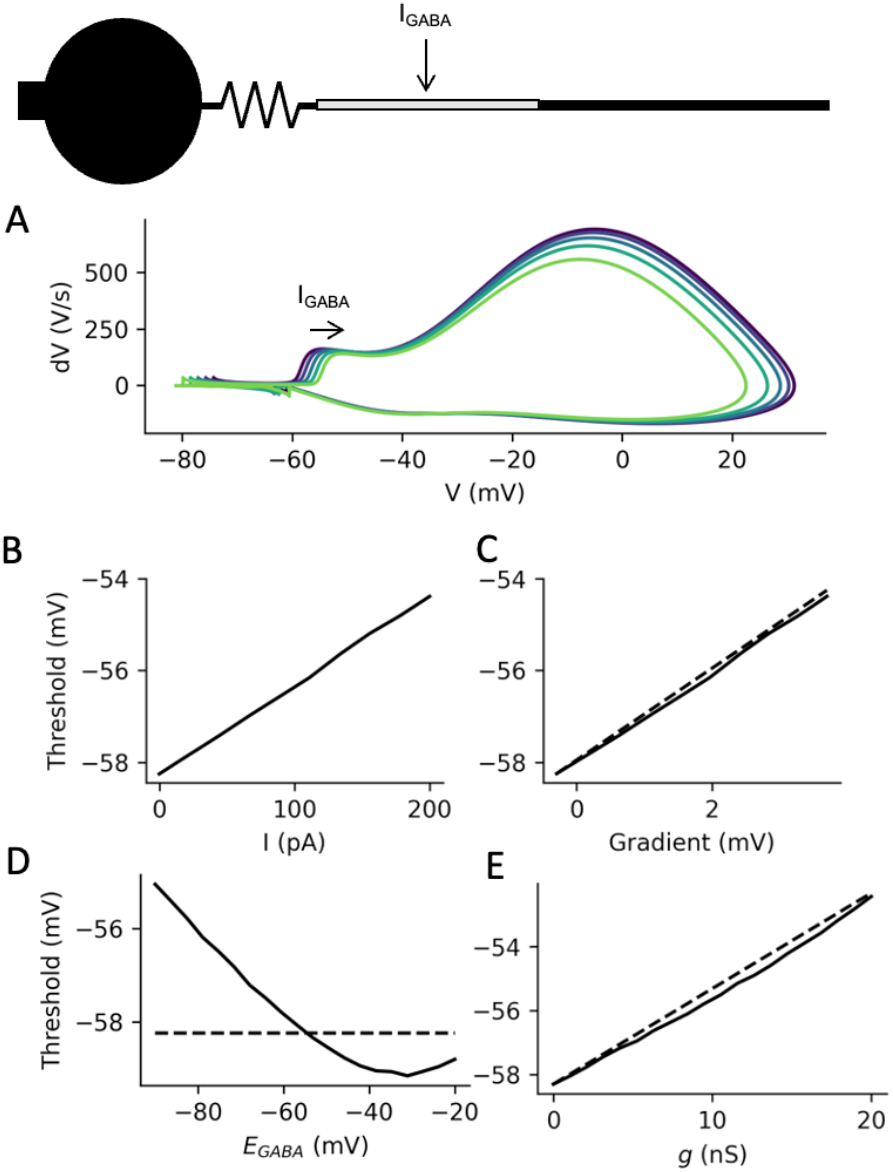
Modulation of action potential initiation by axonal inhibition. A, Phase plots of somatic APs triggered by current pulses at the soma, when a negative current I_GABA_ (0-200 pA) is injected in the middle of the AIS. B, Somatic AP threshold as a function of axonal current. C, Threshold as a function of the mean voltage gradient between soma and AIS, when the axonal current is varied between 0 and 200 pA (dashed: line of slope 1). D, With a synaptic current I_GABA_ = g(E_GABA_-V) (g = 5 nS), threshold as a function of synaptic reversal potential E_GABA_. Dashed line: threshold with no synaptic current at the AIS. E, Threshold as a function of synaptic conductance g (E_GABA_ = -70 mV). Dashed line: theoretical prediction.

Theoretically, the reason is simply that the axonal synaptic current *I* (<0) produces a voltage gradient *R*_*a*_.*I* between soma and current injection site. The AP is triggered when the axonal potential *V* + *R*_*a*_*I* reaches axonal AP threshold V_a_^*^, therefore when the somatic potential *V* reaches 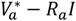. It follows that the AP threshold at the soma is shifted by −*R*_*a*_*I*. This theoretical analysis is based on a point AIS, but it matches results of numerical simulations on a spatially extended AIS, which indeed show an ohmic modulation of somatic AP threshold (Fig. 3B). This modulation precisely matches the difference between somatic potential and average AIS potential, measured before the AP is triggered (Fig. 3C).

As previously mentioned, the synaptic current may be depolarizing or hyperpolarizing, depending on whether the local membrane potential is above or below the reversal potential of chloride: *I = g(E*_*GABA*_ *-V*), where *g* is the synaptic conductance. The effect on AP threshold is therefore determined by the driving force at threshold: *I = g*(*E*_*GABA*_*-V*_*a*_^*^). Thus, in the biophysical model, the axonal synaptic current raises the threshold even when it is depolarizing at rest (Fig. 3D). The threshold modulation reverses when *E*_*GABA*_ is more depolarized than the AP threshold (dashed line). Note that at very depolarized values of E_GABA_, the threshold modulation diverges from linearity: this is because of Nav channel inactivation, which raises the axonal threshold.

*In vitro* measurements indicate that, even when the synaptic current is depolarizing at rest (*E*_*GABA*_ > V_rest_), *E*_*GABA*_ remains about 10 mV below threshold (Woodruff et al., 2009), or is in the same range (Rinetti-Vargas et al., 2017). Substantial Nav channel inactivation would raise the threshold even more. Therefore, the generic effect of this axonal synaptic current should be to raise the somatic AP threshold, at all stages of development.

The synaptic conductance can also increase the axonal AP threshold by opposing the sodium conductance (Goethals and Brette, 2020; Platkiewicz and Brette, 2010), i.e., by “shunting” the AP. It turns out that this effect is approximately equivalent to lowering the synaptic reversal potential by an amount *k*_*a*_, the sodium channel activation slope (about 5 mV), which results in a simple formula for somatic threshold modulation: Δ*V*^*^ = *gR*_*a*_(*E*_*GABA*_ − *V*^*^), where *V*^*^ is the somatic threshold in the absence of axonal synaptic current (see Methods, Theory). Therefore, threshold modulation is predicted to vary linearly with synaptic conductance. This theoretical formula appears to work very well in the biophysical model with spatially extended AIS, where the synaptic current is applied in the middle of the AIS (Fig. 3E).

### Synapse position and structural plasticity of the AIS

How does synapse position along the axon impact AP threshold modulation? Suppose a current *I* is injected on the proximal axon at distance *x* from the soma. As we have seen, this produces an ohmic voltage gradient between soma and current injection site (Fig. 1C). To a first approximation, the axial resistance of a piece of axon is proportional to its length L: *R*_*a*_ = *r*_*a*_*L*, where *r*_*a*_ is the axial resistance per unit length (*r*_*a*_ ≈ 1 MΩ/µm in layer 5 pyramidal cells of the cortex; this value scales with axon diameter *d* as 1/*d*^2^ (Goethals and Brette, 2020)). Therefore, if the AIS is beyond the current injection site, then it gets hyperpolarized by an amount Δ*V*_*AIS*_ = *r*_*a*_*xI*, relative to the soma. If the AIS is between the soma and injection site, then it gets hyperpolarized by an amount Δ*V*_*AIS*_ = *r*_*a*_*x*_*AIS*_*I*. In other words, the effect of an axonal synapse on threshold modulation should increase with its distance from the soma, but only up to the AIS.

This theoretical analysis applies to a point AIS. When we measure threshold modulation as a function of synapse position in a biophysical model with a spatially extended AIS, we find that it increases approximately linearly with synapse position *x* until (approximately) the middle of the AIS, with a slope as predicted from theory, then plateaus (Fig. 4A, 4B). Therefore, synapses are most powerful when placed on the second half of the AIS, and placing them beyond the AIS does not increase their efficiency.

**Figure 4.**
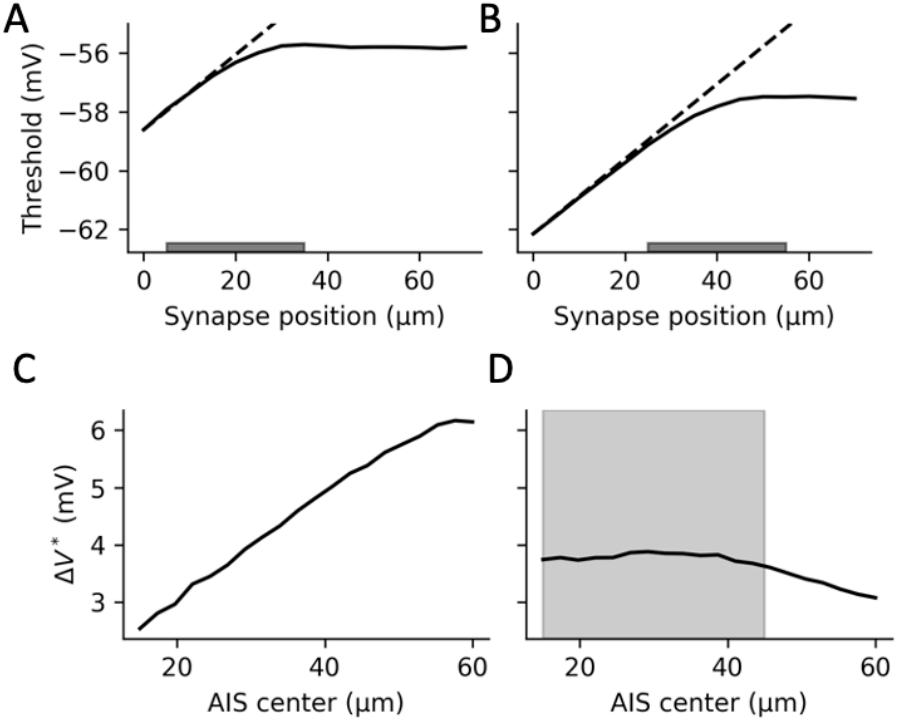
Effect of synapse and AIS position on threshold modulation. A, Threshold as a function of synapse position, when the AIS extends from 5 to 35 µm from the soma (I_GABA_ = -100 pA). Dashed line: theoretical prediction with a point AIS. B, Same as A, but the AIS extending from 25 to 55 µm from the soma. C, Change in threshold (relative to no axonal synaptic current) as a function of AIS center position, when synapses are distributed over the AIS and move with it (E_GABA_ = -90 mV, g = 5 nS). D, Same as C, but synapses are fixed between 15 and 45 µm from the soma (shaded area).

What happens when the AIS moves with activity (Grubb and Burrone, 2010)? If the synapses are placed along the AIS and move together with it, then the impact of those synapses on threshold modulation should increase with distal shifts, as is seen in the biophysical model (Fig. 4C). This change is more modest if synapses are fixed (Fig. 4D).

Empirically, it appears that most axo-axonic synapses are placed on the AIS (Inan et al., 2013; Schneider-Mizell et al., 2021; Somogyi, 1977) and remain in place when the AIS undergoes structural plasticity (Muir and Kittler, 2014; Pan-Vazquez et al., 2020; Wefelmeyer et al., 2015). Our analysis indicates that this ensures a strong effect on AP threshold modulation while preserving some robustness to AIS displacements.

## Discussion

This theoretical analysis can be summarized by the following formula, which quantifies the change in AP threshold at the soma due to a synaptic current on the proximal axon:

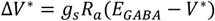

where R_a_ is the axial resistance from the soma to either the synapse or the middle of the AIS, whichever is closer; g_s_ is synaptic conductance, V^*^ is the somatic threshold without a synaptic current. From this analysis, we can conclude that the generic effect of GABAergic currents on the proximal axon is to raise the threshold for AP initiation, even when the current is depolarizing at rest.

This justifies the presumption that axoaxonic cells can potentially exert powerful control on AP initiation. The reason, as the formula emphasizes, is the relative electrical isolation of the AIS provided by the axosomatic coupling resistance *R*_*a*_. There is some uncertainty on the parameter values in the formula, and therefore on the magnitude of the threshold modulation exerted by axoaxonic cells, but it is possible to make some informed estimates. In layer 5 pyramidal cells of the cortex, the middle of the AIS is about 30 µm away from the soma (Hamada et al., 2016), and axonal axial resistance is about 1 MΩ/µm in those neurons (Goethals and Brette, 2020), based on dual soma-bleb patch-clamp recordings (Hu and Bean, 2018). Thus, a reasonable estimate is *R*_*a*_ ≈ 30 MΩ. Tamás and Szabadics (2004) report a peak postsynaptic current of about 50 pA for a single AAC targeting a layer 4 pyramidal cell held at -50 mV, in P24 rats. At that age, *E*_*GABA*_ ≈ −70 mV (Rinetti-Vargas et al., 2017). Therefore, the conductance is *g* ≈ 2.5 nS. Assuming a threshold of *V*^*^ ≈ −55 mV, we obtain Δ*V*^*^ ≈ 1.1 mV for this single spike received from an AAC. If the synaptic conductance does not change with age, then at a later age when *E*_*GABA*_ ≈ −90 mV, we would get Δ*V*^*^ ≈ 2.6 mV.

How would this effect scale *in vivo* when several AACs fire repeatedly on the same AIS? Each AIS receives inputs from ∼4 AACs, for a total of about 20 contacts (Gallo et al., 2020; Inan et al., 2013). Thus, a synchronous discharge from all AACs would raise the threshold by ∼10 mV in an adult. When *n* AACs fire tonically at frequency *F*, the mean synaptic conductance is *nFgτ*_*s*_, where *τ*_*s*_ is the synaptic current decay time. In slices, the latter is about *τ*_*s*_ ≈ 20 ms (Tamás and Szabadics, 2004); a reasonable estimate at physiological temperature might be *τ*_*s*_ ≈ 10 ms. The mean firing rate of AACs during hippocampal theta waves is about 15 Hz (Klausberger et al., 2003). This would make ⟨*g*⟩ ≈ 3.75 nS and therefore ⟨Δ*V*^*^⟩ ≈ 3.9 mV, with a periodic modulation from 0 to 8 mV. Finally, AACs are fast-spiking cells that can fire up to 100 Hz or more (Povysheva et al., 2013). If all AACs on a given AIS fire at maximum rate, this would yield Δ*V*^*^ ≈ 25 mV, essentially blocking AP initiation.

Of course, these are just orders of magnitude with substantial uncertainty. But they show at least that the known physiology and anatomy of AACs are compatible with a strong modulatory effect on AP initiation.

## Materials and Methods

### Theory

Consider a point AIS on the proximal axon, at a distance *x* from the soma. It is electrically connected to the soma through an axial resistance *R*_*a*_ = *r*_*a*_*x*, where *r*_*a*_ is the resistance per unit length (assuming a cylindrical axon). This resistance is related to axon diameter *d* by the formula 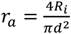, where *R*_*i*_ is intracellular resistivity (of order 100 Ω.cm). For example, with *d* = 1 µm, *r*_*a*_ = 1.3 MΩ/µm.

We then consider a synaptic current *I* = *g*_*s*_(*E*_*GABA*_ – *V*) placed on the AIS, where *g*_*s*_ is the synaptic conductance. The sodium conductance is then opposed by two conductances in parallel: the axial conductance 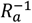 and the synaptic conductance *g*_*s*_. The AP threshold at the AIS varies with the logarithm of the total non-sodium conductance, as *k* 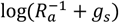, where *k* ≈ 5 mV is the sodium channel activation slope (Platkiewicz and Brette, 2011, 2010). Relative to the absence of synaptic current, the threshold shift is therefore *k* log(1 + *g*_*s*_*R*_*a*_), When *g*_*s*_ is small compared to 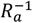 (which is typically the case, see Discussion), this is approximately *kgR*_*a*_ (Taylor expansion). Thus, the axonal AP threshold is 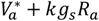, where 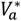 is the axonal threshold without synaptic conductance.

In addition, as this synaptic current flows mostly resistively towards the soma, it introduces a potential difference between soma and AIS equal to −*R*_*a*_*I*_*s*_ = *R*_*a*_*g*_*s*_(*V*_*a*_ − *E*_*GABA*_). Thus, an AP is initiated when 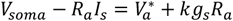. Thus, the AP threshold at the soma is:

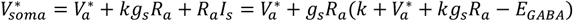

Therefore, assuming again that *g*_*s*_*R*_*a*_ is small, the shift in somatic threshold relative to an absence of synaptic conductance is:

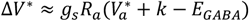

As it turns out, the somatic threshold in the absence of synaptic conductance 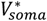 is depolarized relative to the axonal threshold, by an amount *k* (≈ 5 mV) (Brette, 2013), in agreement with measurements (Kole and Stuart, 2008). Therefore, the final formula is simply:

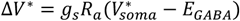

The theory with a spatially extended AIS is more complex, but we previously found that the threshold in an extended AIS is close to the value obtained in a point AIS with the same total conductances, placed at the middle of the extended AIS (Goethals and Brette, 2020).

### Numerical simulations

Models were simulated with the Brian2 simulator (Stimberg et al., 2019), using a 1 µs timestep. The biophysical model is described in (Goethals and Brette, 2020). Briefly, it consists of a 500 µm long axon (diameter 1 µm), a 30 µm soma and a 1 mm long dendrite (diameter 6 µm) with voltage-gated sodium and potassium channels. Parameter values are listed in Table 1.

**Table 1.** Parameters values of the biophysical model. Time constants corrected for temperature are indicated in brackets.

|  |  |  |
| --- | --- | --- |
| Passive properties | $R_m$ | 15 000 $\Omega \cdot \text{cm}^2$ |
| | $E_L$ | -75 mV |
| | $R_i$ | 100 $\Omega \cdot \text{cm}$ |
| | $C_m$ | 0.9 $\mu\text{F}/\text{cm}^2$ |
| Nav channels | $g_{Na, \text{soma}}$ | 250 S/m<sup>2</sup> |
| | $g_{Na, \text{dendrite and axon (non AIS)}}$ | 50 S/m<sup>2</sup> |
| | $g_{Na, \text{AIS}}$ | 4000 S/m<sup>2</sup> |
| | $E_{Na}$ | 70 mV |
| | $V_m^{1/2}, \text{soma}$ | -30 mV |
| | $V_h^{1/2}, \text{soma}$ | -55 mV |
| | $V_m^{1/2}, \text{AIS}$ | -35 mV |
| | $V_h^{1/2}, \text{AIS}$ | -60 mV |
| | $k_m$ | 5 mV |
| | $k_h$ | 5 mV |
| | $\tau_m^*$ | 150 μs (corrected: 54 μs) |
| | $\tau_h^*$ | 5 ms (corrected: 1.8 ms) |
| Kv1 channels | $g_K, \text{soma}$ | 250 S/m<sup>2</sup> |
| | $g_K, \text{dendrite and axon (non AIS)}}$ | 50 S/m<sup>2</sup> |
| | $g_K, \text{AIS}$ | 1500 S/m<sup>2</sup> |
| | $E_K$ | -90 mV |
| | $V_n^{1/2}$ | -70 mV |
| | $k_n$ | 20 mV |
| | $\tau_n^*$ | 1 ms |

AP threshold is measured as the inflexion point of the phase plot (maximum of dV^2^/dt^2^), which is similar to a threshold on dV/dt but more robust (Telenczuk et al., 2017).

## Code availability

Code can be found at https://github.com/romainbrette/theory-of-axoaxonic-inhibition.

## Acknowledgments

We thank Marcel Stimberg for technical assistance.

This work was supported by Agence Nationale de la Recherche (ANR-20-CE30-0025-01, ANR-21-CE16-0013-02 and ANR-23-CE16-0020-02).

